# Transcriptional profiling of potato (*Solanum tuberosum* L.) during a compatible interaction with the root-knot nematode, *Meloidogyne javanica*

**DOI:** 10.1101/849414

**Authors:** Teresia Nyambura Macharia, Daniel Bellieny-Rabelo, Lucy Novungayo Moleleki

**Author notes:** Correspondence: Prof. Lucy Moleleki.

## Abstract

Root-knot nematode (RKN, *Meloidogyne javanica*) presents a great challenge to *Solanaceae* crops, including the potato. In this report, we conducted an investigation to understand the transcriptional regulation of molecular responses in potato roots during a compatible interaction following RKN infection. In this study, analysis of gene expression profiles using RNA-seq of *Solanum tuberosum* cv Mondial with RKN interaction at 0, 3- and 7-days post-inoculation (dpi). In total, 4,948 and 4,484 genes were respectively detected as differentially expressed genes (DEGs) at 3 and 7 dpi. Functional annotation revealed that genes associated with metabolic process were enriched at the transcriptional level suggesting they have an important role in RKN disease development. Nematode infection caused down-regulation of 282 genes associated with pathogen perception hence interfering with activation plant immune system. Further, late activation of pathogenesis-related genes, down-regulation disease resistance genes and activation of host antioxidant system contributed to a susceptible response. Activation of Jasmonic acid (JA) pathway and protease inhibitors was due to wounding during nematode migration and feeding. Nematode infection suppressed ethylene (ET) and salicylic acid (SA) signalling pathway hindering SA/ET responsive genes involved with defense. Induction of auxin biosynthesis genes, regulation of cytokinin levels and up-regulation of transporter genes facilitated of nematode feeding sites (NFSs) initiation. The regulation of several families of transcription factors (TFs) in the plant, such as WRKY, GRAS, ERF BHLH and MYB, was affected by RKN infection disrupting plant defense signalling pathways. This clearly suggest that TFs played an indispensable role in physiological adaptation for successful RKN disease development. This genome-wide analysis revealed the molecular regulatory networks in potato roots which are successfully manipulated by RKN. Being the first study analysing transcriptome profiling of RKN diseased potato, it will provide unparalleled insight into the mechanism underlying disease development.

## Introduction

Potato, *Solanum tuberosum* (L) belongs to the *Solanaceae* family, which comprises several economically important crops such as tomato, pepper, aubergine, and tobacco. Plant parasitic nematodes particularly root-knot nematodes, are among the most destructive and economically important pests of potatoes worldwide (Scurrah et al., 2005, Jones et al., 2017). In this context, *Meloidogyne* spp are obligate and highly polyphagous pests that form an intricate relationship with their host causing drastic morphological and physiological changes in plant cell architecture (Gheysen & Fenoll, 2002). A typical life cycle of RKNs spans between 4-6 weeks depending on the nematode species and environmental conditions. Following the embryonic phase, the infective second-stage juveniles (J2s) hatch from the egg. At 3dpi the nematodes have already penetrated the host root tips and migrating towards the elongation zone (Castaneda et al., 2017). At this stage the J2s select target cells to initiate reprogramming of host cells to giant cells (GCs). The nematodes are completely dependent on the induced GCs for supply of nutrients. During the induction stage the parasitic J2 abandons its migratory lifestyle becomes sedentary to concentrate on feeding, development and reproduction (Bartlem et al., 2013). As the GCs enlarge, surrounding cells undergo rapid division causing swelling of roots and discontinuity of the vascular tissue. The sedentary nematode further moults into J3, J4 stages and finally into the adult stage when the nutrient acquisition stage commence from 7dpi. The developing, nematode, GCs and surrounding tissue contribute to the formation of RKN symptom (Bartlem et al., 2013). Analogous to other plant pathogens, nematode secretions play a crucial role in manipulation of the host cellular function. Secreted molecules suppress host defense to initiate a successful infection process including establishment and maintenance of NFSs (Hewezi & Baum, 2013). In the genus *Meloidogyne*, several effectors have been reported such as: MiLSE5, which interferes with host metabolic and signalling pathways; or MjTTL5, Misp12 and MgGPP, which suppress the host immune responses facilitating successful nematode parasitism (Vieira & Gleason, 2019).

Due to their capability to infect plant species from diverse families, RKN species pose a great challenge to crop production globally (Sasser & Freckman, 1987). In 2014, 22 species of RKN were reported in Africa causing damage to various vegetable and field crops (Onkendi et al., 2014). Both tropical and temperate RKN species are present in potato growing regions of South Africa with *M. javanica* and *M.incognita* being the prevalent species impairing the potato production sector (Onkendi & Moleleki, 2013). For decades the use of nematicides has been effective in managing RKN populations. However, their usage is coupled with adverse effects to the ecosystem. This has led to withdrawal of the most effective nematicides from the agro-markets, further aggravating crop losses due to RKN (Onkendi et al., 2014). Plant host resistance through the use of resistant cultivars is an effective and environmentally safe alternative method of controlling RKN species (Onkendi et al., 2014). Nevertheless, the current cultivated potato cultivar lack resistance against RKN (Dinh et al., 2015). Thus, studies involving plant-nematode interactions will deepen our understanding of the molecular regulatory networks associated with resistance or susceptibility. The insights drawn from such studies will be useful in breeding programs to develop novel target-specific control strategies against nematodes.

RNA-Seq has become a powerful instrument for gene expression profiling and detection of novel genes (Wang et al., 2009, Ozsolak & Milos, 2011) which has been used widely to study the expression profiles of RKN diseased Solanaceae plants (Xing et al., 2017, Li et al., 2018, Shukla et al., 2018). RNA-seq profiling has been used to decipher potato responses to various abiotic (Zuluaga et al., 2015, Gálvez et al., 2016) and biotic stresses (Kwenda et al., 2016, Yang et al., 2018) where large sets of genes and pathways associated with either biotic and abiotic stress were revealed. To date most research has focused on potato gene expression in response to potato cyst nematodes (Jolivet et al., 2007, Walter et al., 2018, Kooliyottil et al., 2019) while potato responses to RKN infection remain poorly understood. Here, we set out to evaluate the responses of potato cultivars to root-knot nematode infection. Our results revealed that seven commercially tested South African potato cultivars were susceptible to *M. javanica.* Further, in order to investigate the molecular basis of this compatible interaction, we employed RNA-Seq to analyse differential gene expression patterns in *Solanum tuberosum* cv. Mondial subjected to *M. javanica* infection at two early stages (3 and 7dpi).

## Results and Discussion

### Susceptibility of potato cultivars to *Meloidogyne javanica*

In this study, we evaluated the susceptibility to RKN in seven commercially available potato cultivars in South Africa. The number of galls induced, and reproductive potential of the nematodes was used to assess the host status of the potato cultivars. Our results show that all the seven potato cultivars were efficient hosts to *M. javanica* as indicated by the high reproductive factor (Rf >1) **(Fig 1A).** This further supports the findings by Pofu and Mashela (2017) which concluded that South African potato cultivars are efficient hosts to *Meloidogyne* species. Based on the gall numbers, cultivars were classified as highly susceptible (BP1, Mondial and Lanorma), susceptible (Up-to-date, Sifra, and Valor), and moderately resistant (Innovator) **(Fig 1B**) according to the ranking scale coined by Taylor and Sasser (1978) *Meloidogyne Javanica* infection generally reduces plant growth and yield of potato (Vovlas et al., 2005). Similarly, RKN infection caused reduction in root length and shoot length in various potato cultivars compared to their corresponding controls (**Fig 1 C and D**). The reduced growth is attributed to root injury caused during nematode penetration and feeding, which impairs the plant root systems, hence reducing the efficiency of roots to absorb water and nutrients. As a result, the top growth of the plant is reduced, and this explains the reduced shoot lengths. Interestingly, increased root length was recorded in four diseased cultivars in comparison to untreated controls. The production of secondary roots may be a characteristic of nematode infected plant’s efforts to recompense for root injury (Mcdonald & Nicol, 2005). In addition, mature galls exhibited either a single or more egg masses, which indicates that *M. javanica* was infective on potato cultivars (**Fig 1E and F)**. Therefore, nematodes were able to penetrate the host system, subdue the host defense responses during the entire infection process, complete their life cycle and reproduce.

**Figure 1:**
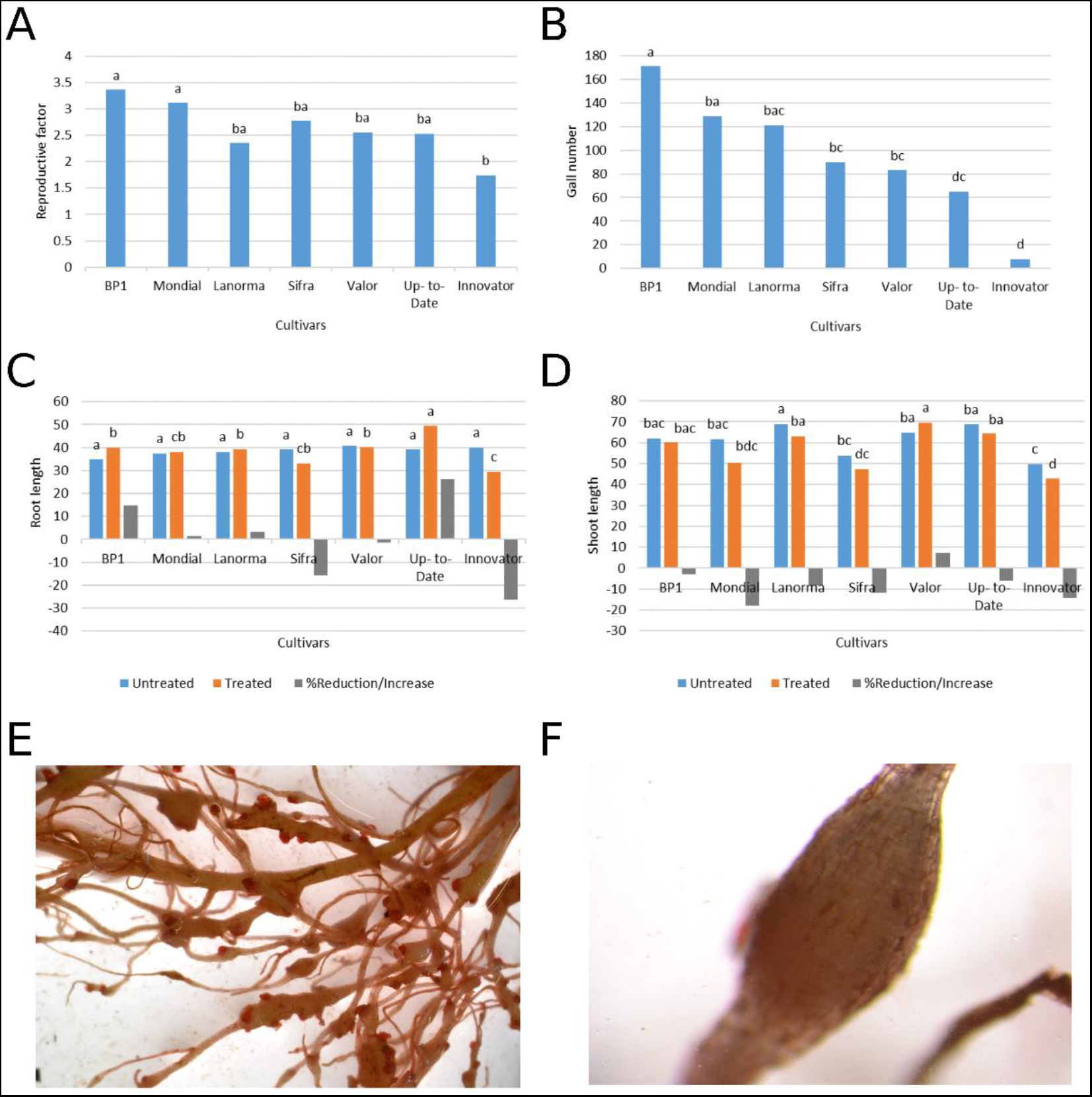
Responses of seven potato cultivars to *M. javanica* infection. **(A)** and **(B)** Reproductive factor and the number of galls, respectively induced by RKN. **(C)** and **(D)** Show the effect of nematode infection on root length and shoot length of potato cultivars. % increase or reduction = 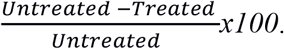 Values are means of five replicates. Statistical significance between the cultivars was determined by one-way ANOVA analysis with Fisher’s least significant difference test at P< 0.05. (E) and (F) show nematode damage on potato roots, the egg masses stained pink and a mature gall, respectively.

### Transcriptome data analysis and functional annotation of differentially expressed genes

Aiming to understand the molecular basis of this compatible interaction between RKN and potato, RNA sequencing was performed on the highly susceptible cultivar *Solunum tuberosum* cv Mondial. Two infection stages were selected for the analysis: 3 and 7 dpi. These time points correspond to nematode stages of induction of feeding sites at 3 dpi and nutrient acquisition stage that starts from 7 dpi to 8 weeks after infection (Bartlem et al., 2013). Approximately 1.3 billion paired-end reads were generated yielding an average of 23 million high quality reads for individual samples. Successfully mapped reads onto the *S. tuberosum* reference genome (v4.03) (Consortium, 2011) accounted for 78-86% of the total of reads generated per sample (**Table S1**). Log2 fold change ≥ ± 1 and adjusted p-value (FDR) < 0.05 were used as cut off values to obtain DEGs through pairwise comparison between the mock-inoculated samples and infected samples at 3 and 7 dpi. Overall, 4948 potato genes were differentially expressed at 3dpi. Of these, 2867 were down-regulated and 2081 up-regulated. At 7dpi, 2871 and 1613 genes were detected to be down and up-regulated respectively **(Fig 2A)**. Collectively, 3108 genes were regulated at both 3 and 7 dpi: 2069 down- and 1022 up-regulated. As biotrophic organisms, RKNs need to actively suppress the host defense during the infection process. This might explain the current observation where 57.75% of the DEGs (3,652 out of 6,324) were suppressed (**Fig 2A and B**).

**Figure 2:**
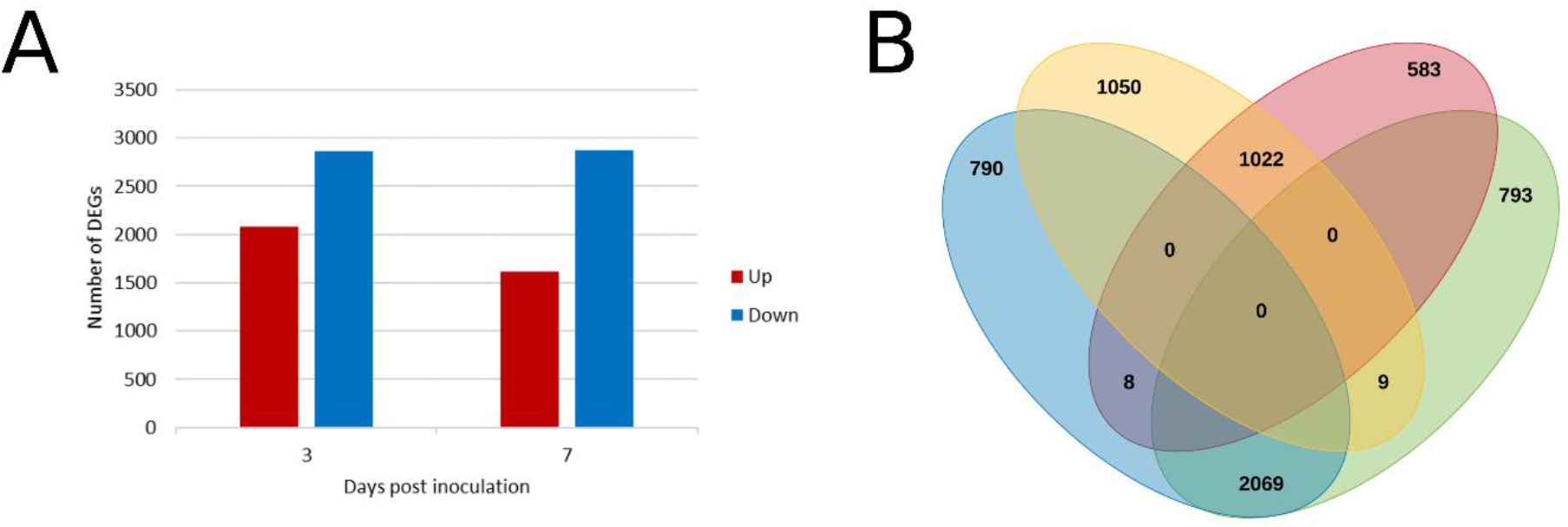
Schematic representation of DEGs in potato roots following *M. javanica* infection **(A)** Number of differentially expressed genes detected at 3 and 7 dpi compared to the mock-inoculated samples. ‘Down’ designates down-regulated genes. ‘Up’ designates up-regulated genes. **(B)** Venn diagram of the distribution of DEGs between 3 and 7 dpi. Yellow and blue ovals represent up-regulated and down-regulated DEGs at 3dpi, respectively. Red and green ovals indicate the genes upregulated and downregulated at 7 dpi, respectively.

GO enrichment analysis was performed using the AgriGo tool (Tian et al., 2017) to reveal the main regulatory trends in root tissues on the course of RKN infection. The GO terms were grouped into three main functional categories at adjusted *p-value < 0.05* and categorized using WEGO software (Ye et al., 2018). Within the biological process class, the highest percentage of the DEGs was down-regulated and fell under metabolic process category. Within this category, we found the following sub-categories: Primary metabolic process, cellular metabolic process, biosynthetic process, oxidation-reduction process and regulation of metabolic process. Accordingly, past studies revealed that nematode infection modulates the expression of genes involved in metabolic activities, particularly the primary metabolic process due to the high demand for nutrients and energy (Hofmann et al., 2010). Other significant GO terms in this category include response to stimulus, cellular process, localization and signalling processes and regulation of biological processes **(Fig 3 and S2 Table).**

**Figure 3:**
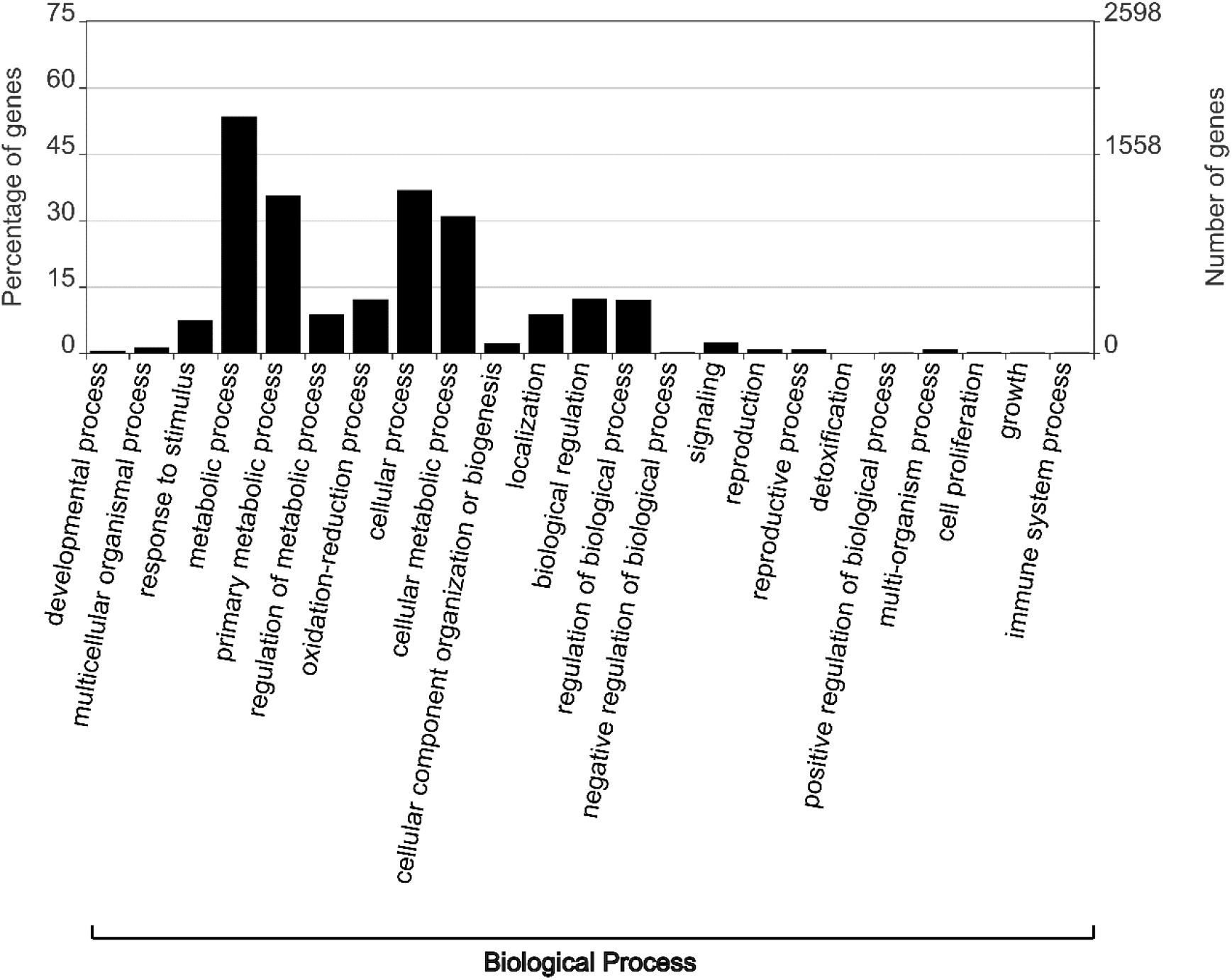
A representation of GO analysis demonstrates the percentage of DEGs enriched within the Biological Process category.

### Plant signal transduction, pathogen perception, and defense-related genes are modulated by *M. javanica* infection

Plants have developed the innate immune system to inhibit pathogen invasion and multiplication. Pattern triggered immunity (PTI) (plants first line of defence) relies on perception of pathogen/damage-associated molecular patterns (PAMPs/DAMPs) by pattern recognition receptors (PRRs) (Zipfel, 2014, Macho & Zipfel, 2014). Plant PRRs are either surface localized receptor-like proteins (RLPs) or receptor-like kinases (RLKs) that perceive and transmit danger signals to activate defense response (Zipfel, 2014). In this study, genes encoding for RLKs and RLPs (e.g. serine-threonine protein kinase, leucine-rich repeat (LRR) receptor-like protein kinase) and wall-associated receptor kinases (WAKs) were detected among the DEGs. The majority of the PRRs (68.28%) were repressed by nematode infection at 3 and/or 7 dpi following nematode infection **(Fig 4A and S3 Table)**. Previous reports show that phytonematodes are able to induce plant basal defense responses through recognition by large arsenals of plant receptors (Peng & Kaloshian, 2014, Teixeira et al., 2016, Mendy et al., 2017). Mendy et al. (2017) reported the first surface localized LRR receptor-like kinase (NILR1) that was up-regulated in response to nematode attack in *Arabidopsis.* Arabidopsis *nilr1* mutants were found to be hyper susceptible to a wide range of phytonematodes (Mendy et al., 2017). Similarly, our results revealed that six out of seven WAK-encoding genes were down-regulated by nematode infection **(Fig 4A and S3 Table).** The WAK proteins perceive danger signals to activate PTI responses (Ferrari et al., 2013). Past research shows that WAK proteins are important components of potato disease resistance against various microbes (Kwenda et al., 2016, Yang et al., 2018). For instance, WAK genes were induced in a tolerant cultivar (BP1) which correlated with enhanced perception of *Pectobacterium brasiliense* (formerly *Pectobacterium carotovorum* subsp *brasiliense*) (Kwenda et al., 2016). Similarly, WAK genes were up-regulated conferring resistance to *Phytophthora infestans* in potato genotype SD20 (Yang et al., 2018) Additionally, plants defective of PRRs or PTI signaling components are typically susceptible to both adapted and non-adapted microbes (Macho & Zipfel, 2014). This notion is confirmed by *M. javanica* ability to interfere with the functioning of RLKs, RLPs and WAK genes lead to successful disease development.

**Figure 4:**
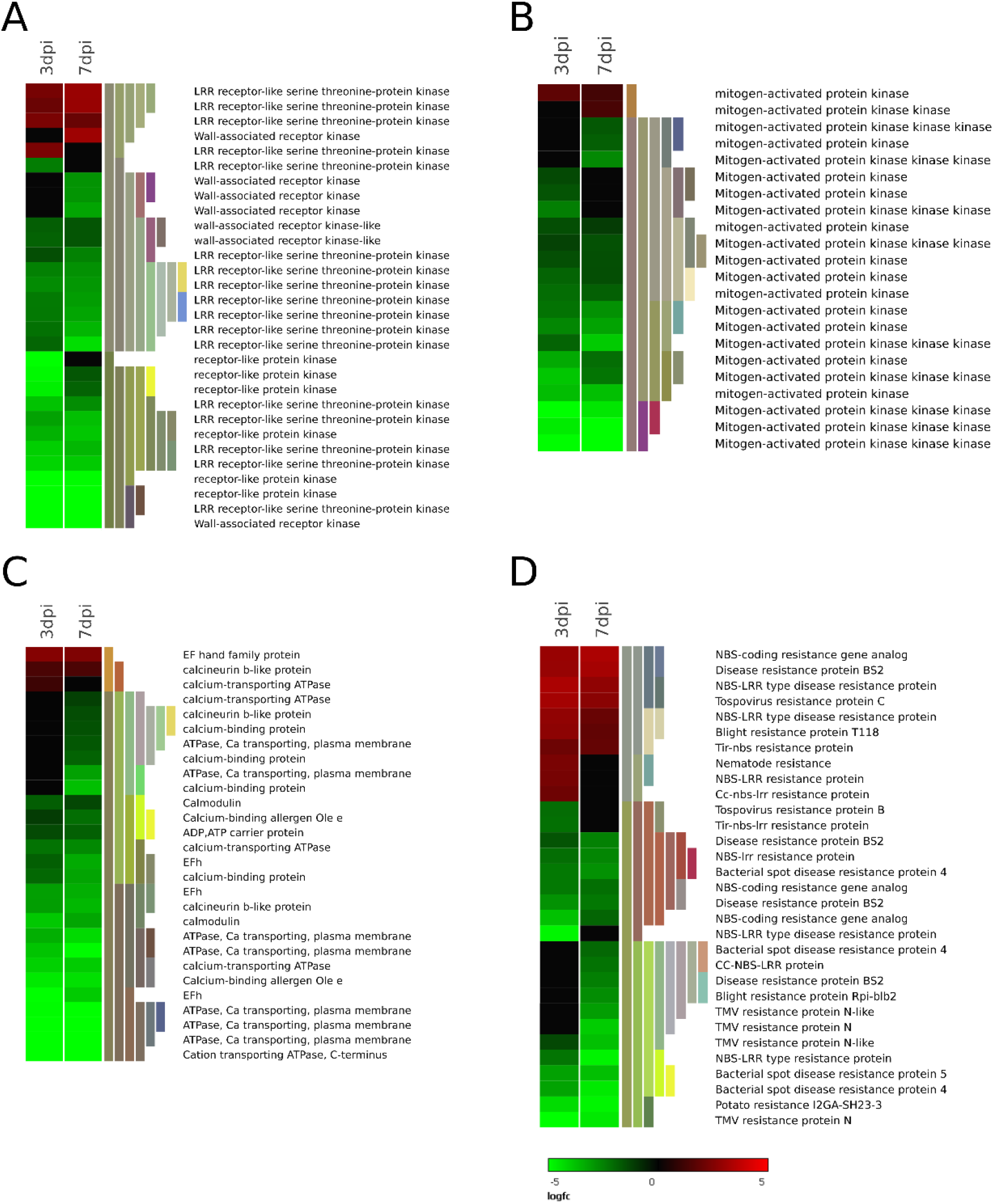
Heat map representation of selected DEGs associated with pathogen perception **(A)** PRRs-RLKs, RLPs and WAKs **(B)** MAPK signaling pathways **(C)** Ca2+ signaling pathways, and Disease resistance proteins. **(D)** (The heat map illustrates a subset of genes from each group. Refer supporting information for all DEGs in each group).

Transmission of perceived signals from the PRRs is mediated through the MAPK cascade and calcium (Ca^2+^) signaling pathway which transfers downstream components of plant immunity. Here, we found the expression of MAPKs genes was largely repressed (20 out of 22 genes) by nematode infection **(Fig 4B and S3 Table)**. The MAPK cascade basically entails three-tiered kinases (a) a MAP kinase kinase kinase (MAPKKK) (b) a MAP kinase kinase (MAPKK), and (c) the MAP kinase (MAPK) which mediates transmission of extracellular signals to activate an appropriate defense output (Jagodzik et al., 2018). The role of MAPK in plant defense against nematodes has been demonstrated previously. It was reported by Zhang et al. (2017) and Postnikova et al. (2015) that the induction of MAPK genes leads to resistance against cyst nematode and RKN, respectively in soybean plants. Further, *M. javanica* repressed 91.6% of the genes involved in Ca^2+^ signaling pathway (33 out of 36) **(Fig 4C and S3Table)**. This includes calmodulin (CaM), calcineurin B-like proteins (CBL), Ca^2+^ dependent kinases (CPKs) and Ca^2+^ receptors that transmit Ca^2+^ signatures into a specific cellular and physiological response after a pathogen challenge (Zhang et al., 2014). Takabatake et al. (2007) demonstrated that repression of CaM/CML members’ expression or loss of function in mutated plants strongly affects immunity. Furthermore, in plant-nematode interaction, calcium burst was associated with the release of ROS causing cell death and inhibiting establishment of (GCs) in potato (Davies et al., 2015). In connection to this, our results indicate that *M. javanica* ability to repress MAPK and Ca^2+^ pathways interfered with the transmission of signals responsible for activation of precise and prompt defense response, hence a susceptible response.

NBS-LRR disease resistance proteins have been implicated in mediating resistance against various phytonematodes (Williamson & Kumar, 2006). In our study, 114 disease resistance genes (out of the 641 in the potato genome) (Sharma et al., 2013) were differentially expressed following root knot nematode infection. The highest proportion of these proteins (54.35%) including NBS-LRR disease resistance proteins, was repressed at 3 and/or 7 dpi **(Fig 4D and S3 Table).** Similarly, repression of 12 NBS-LRR genes by cereal cyst nematode in wheat led to a susceptible response (Qiao et al., 2019). Altogether, down-regulation of resistance genes indicates repression of plant resistance by *M. javanica* infection.

Apart from the activation of MAPK and Ca^2+^ signaling, PTI activation is associated with expression of pathogenesis related (PR) proteins. In this study, we found the expression of several PR proteins under the regulation of *M. javanica* including 11 chitinase encoding genes (PR-3 and PR-4). The PR-3 and PR-4 are markers for JA-mediated defense response with 8 genes specifically up-regulated at 7dpi **(Fig 5A and S3 Table).** In a similar fashion, it has been previously reported that increased chitinase activity do not correlate with resistance in potato to potato cyst nematode (Wright et al., 1998). Therefore, chitinase might be functioning as a signalling molecule to stimulate other PR proteins, or alternatively, its induction may be due to wounding response (Wright et al., 1998). Regarding SA-responsive genes (PR-1 and PR-5), 14 were up-regulated and 11 were down-regulated by nematode infection specifically at 7dpi **(Fig 5A and S3 Table).** According to previous studies, the PR-5 transcripts were induced following RKN and CN infection in *S. lycopersicum* and *Brassica nigra*, respectively (Sanz-Alférez et al., 2008, van Dam et al., 2018). These observations are corroborated in the current study where the majority of thaumatin-like and osmotin genes (55.82%) were induced by nematode infection. Moreover, the stimulation of PR-5 protein has been associated with osmotic stress induced during nematode invasion (Sanz-Alférez et al., 2008). Generally, there was delayed activation of the PR genes as the majority of the PR encoding genes were induced at 7 dpi. This could reflect the strategy adopted by the RKN to suppress PR encoding genes in early stages of colonization to ensure successful nematode infection.

**Figure 5:**
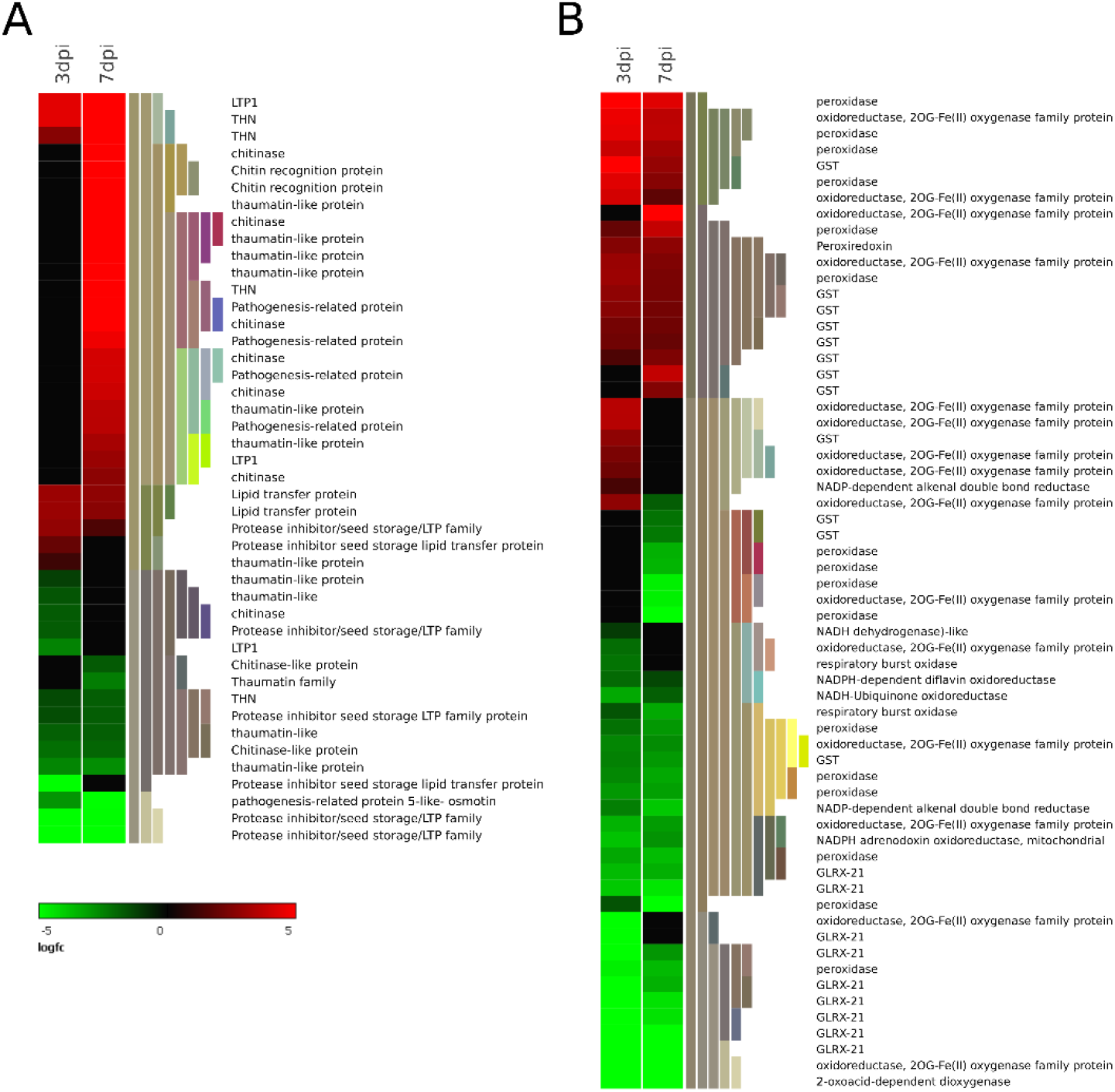
Heat map representation of selected DEGs associated with plant defense **(A)** Pathogenesis-related protein. **(B)** Oxidative stress-related gene (The heat map illustrates a subset of genes from each group. Refer to supporting information for all DEGs in each group).

Rapid generation of reactive oxygen species (ROS) is one of the early PTI cellular events that trigger a number of defense responses such as activation of several defense genes and cell wall reinforcement (Goverse & Smant, 2014) In this study, NADPH oxidase, respiratory burst homologue (RBOHs) and peroxidases, which are important players in production of ROS in plants, were differentially regulated both at 3 and 7dpi **(Fig 5B and S3 Table)**. Genes encoding for peroxidases (30 genes) were repressed to a larger extent at 7 dpi than at 3 dpi **(Fig 5B and S3 Table)**. This implies that *M. javanica* suppresses ROS-mediated defense signaling during induction and acquisition of nutrients in the GCs. In addition, two genes coding for 2-oxoacid-dependent dioxygenase were down-regulated at 3 and/or 7 dpi. 2-oxoacid-dependent dioxygenase enzyme mediates a variety of oxidative reactions and synthesis of secondary metabolites (Prescott & Lloyd, 2000) and has toxic effects on a wide range of pathogens including phytonematodes (Hansen et al., 2008). Genes encoding for 2OG-Fe (II) oxygenase superfamily were up-regulated (23 genes out of 34) following nematode infection in this study **(Fig 5B and S3 Table)**. Patel et al. (2010) reported that the interaction between host 2OG-Fe (II) oxygenase and a nematode effector HS4F01 increased the plant susceptibility to cyst nematode. This illustrates that *M. javanica* effectors might interact with the host proteins responsible for oxidative responses hence interfering with ROS mediated defense signaling.

Genes encoding for glutathione, glutaredoxin, thioredoxin, peroxiredoxins, ascorbate and peroxidases comprise plant’s antioxidant network, which is responsible for controlling ROS levels (Laporte et al., 2012). These genes were differentially regulated by nematode infection in this study **(Fig 5B and S3 Table)**. Emerging evidence shows that RKN can utilize the host ROS scavenging system to reduce the damaging effects of oxygen species (Lin et al., 2016, Guan et al., 2017). Here, we detected one gene encoding for peroxiredoxin (PGSC0003DMG401002721), the main detoxifying antioxidant enzyme in the plant-nematode interface (Goverse & Smant, 2014), being up-regulated at both timepoints **(Fig 5B and S3Table)**. In addition, 16 out 23 genes encoding for glutathione S transferase (GST) and UDP-Glycosyltransferase (6 out of 9 genes) **(S3 Table)** were up-regulated following *M. javanica* infection **(Fig 5B and S3Table)**. Qiao et al. (2019) reported that CN nematode can utilize the GST and UDP-Glycosyltransferase antioxidant enzymes to ameliorate the ROS effects as well control plant defense. In this context, it is likely that in the current interaction, the nematode activated host antioxidant mechanism to interfere with defense response and to avoid the harmful effect of ROS molecules.

### Nematode responsive transcription factors

Several transcription factors (TFs) were detected as DEGs in response to *M. javanica* infection. This includes ERF (77), MYB and MYB-related (62), bHLH (49), bZIP (23), WRKY (33) and GRAS (32). In total, these differentially expressed TFs represent 75% of the TFs found in *S. tuberosum*. Most of the differentially expressed TFs in the current data set were down-regulated (298/532) after nematode infection **(Fig 6A and S4 Table)**. Classification and identification of the differentially expressed TFs was attained from the Plant Transcription Factor Database (http://planttfdb.cbi.pku.edu.cn/ v .4.0) (Jin et al., 2016).Transcription factors (TFs) are key regulators of plant response to various biotic stress in potato. For instance, in response to *P. brasiliense* infection in potato, 4 families of TFs (WRKY, bHLH, MYB, and AP2/ERF) were regulated (Kwenda et al., 2016). Similarly, in potato, several TFs were found to be important regulators of resistance response against *P. infestans* (Yang et al., 2018).

**Figure 6:**
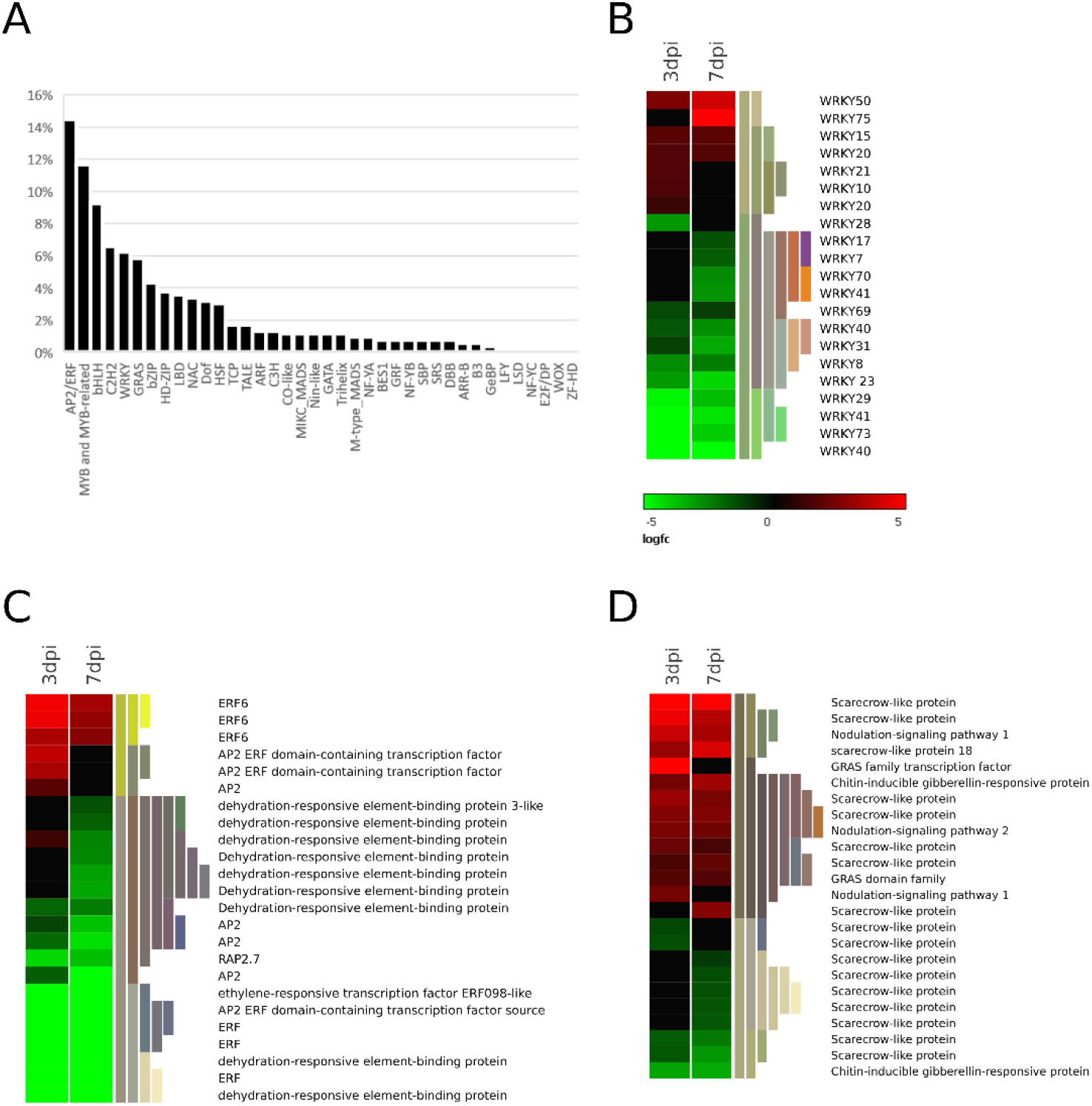
Heat map representation of differential regulation of TFs. **(A)** Represents various families of TFs under the regulation of RKN. **(B)** WRKY family. **(C)** AP2/ERF family. **(D)** GRAS family. (The heat map represents a subset of the differentially expressed family of TFs. Refer to supporting information for all TFs family displaying differential expression).

The ERF TFs are associated with hormone signal transduction of sacylic acid (SA), jasmonic acid (JA) ethylene (ET), and PR via binding to the GCC box of target genes that positively or negatively regulate transcription of various stress responses (Li et al., 2017). In this study, most of the genes encoding for AP2/ERF TF family were down-regulated (70.37%) at 3 and/or 7 dpi. In addition, 15 genes were suppressed to a greater extent at 7 than 3 dpi **(Fig 5C and S4 Table)**. This could be ascribed to the secretion of nematode effectors and subsequent suppression of defense response associated with the activation of AP2/ERF TFs. Among the down-regulated ERF TFs we found 7 genes encoding DREB (out of 9 DREB genes), which are regarded as main regulators of abiotic stress responses (Zhou et al., 2010). **(Fig 6C and S4 Table)**. Qiao et al. (2019) recently reported that two genes encoding for DREB were strongly repressed in wheat following a compatible interaction with CN. This indicates that DREB genes may regulate signaling pathways associated with defense response to nematode infection. This can be subjected to further analysis to identify their specific role in plant-nematode interactions. Further, we found three ERF6 TFs activated in response to *M. javanica* infection **(Fig 6C and S4 Table)**. In this context, ERF6 has been described to positively regulate JA/ET and resistance against *Botrytis cinerea* in *A. thaliana* (Moffat et al., 2012). In addition, Warmerdam et al. (2019) showed that ERF6 regulates *M. incognita* disease development in *A. thaliana.* ERF6-mutated plants recorded a higher number of RKN egg masses indicating a role of ERF6 in enhancing host susceptibility to *M. incognita* as a result of deteriorated plant defenses (Warmerdam et al., 2019). In this case, ERF6 TFs have a role in mediating potato susceptibility to RKN. Our findings indicate that down-regulation of AP2/ERF TFs following nematode infection could have debilitated plant defense through targeting the defense signaling pathways regulated by the AP2/ERF family of TFs.

It is generally accepted that pathogen-directed modulation of WRKY genes in plants is an important aspect that enhances success rates of pathogen infection. Cyst nematode’s successful infection process in *A. thaliana* roots was attributed to the nematode’s control over the expression of WRKY genes (Ali et al., 2014). In agreement with that notion, we found 23 genes down-regulated WRKY-encoding genes, including *WRKY40, WRKY23,* and *WRKY29* at both infection stages **(Fig 6B and S4 Table)**. In cotton plants*, GhWRKY40* has been reported to regulate wounding and resistance response against *Ralstonia solanacearum* (Wang et al., 2014). Furthermore, the up-regulation of *WRKY23* influenced an early resistance response to *M. incognita* infection in cucumber plants (Ling et al., 2017). It has also been reported that in *Arabidopsis, AtWRKY29* is an important constituent of MAPK mediated defense pathway against microbes (Asai et al., 2002). Here we detected 20 genes encoding for MAPK suppressed by nematode infection. The suppression of *WRKY29* might have influenced the expression of *MAPK* genes interfering with transmission of signals that elicit a defense response. Moreover, we found WRKY75 to be up-regulated by nematode infection at 7dpi. In tomato, *SlyWRKY75* regulates the JA-signal transduction system (López-Galiano et al., 2018) indicating activation of the JA pathway by *M. javanica* infection. Our data reveals that RKN interferes with important defense signaling components such as MAPK and JA pathways eliciting a susceptible response through down regulation of WRKY TFs.

Among the 34 down-regulated MYB TFs **(S4 Table)**, we found three genes encoding for MYB108 at 3 and 7 dpi. *Arabidopsis* AtMYB108 has been characterized as an important regulator of both biotic and abiotic stresses (Mengiste et al., 2003) It is also known that the expression of *GhMYB108* in cotton, responds to application of defense-related phytohormones such as SA, JA and ET (Cheng et al., 2016). The absence of *GhMYB108* led to increased susceptibility of cotton plants to *Verticillium dahliae* infection while its ectopic overexpression enhanced tolerance to the fungal pathogen (Cheng et al., 2016). This would, therefore, indicate that down-regulation of MYB108 coding genes interfered with the defense signaling pathway leading to a compatible response. MYC2, MYC3, and MYC4 from the bHLH family are a part of the JA signal transduction system (Pireyre & Burow, 2015). bHLH activates various sets of plant genes in response to environmental factors such as phytohormone signaling, and development (Pireyre & Burow, 2015). Here, we found 49 genes encoding for bHLH TFs with a total of 23 up-regulated and 24 down-regulated in response to RKN infection **(S4 Table)**. Our results indicate that nematode infection interferes with these important regulators of JA mediated defenses by blocking the expression of some of the bHLH TFs responsible for mounting sufficient defense responses against *M. javanica*.

Out of 71 GRAS representatives in the potato genome, 32 were differentially expressed following nematode infection in the present data set, out of which 65.63% were up-regulated **(Fig 6D and S4 Table)**. This includes 17 scarecrow-like (SCL) encoding genes important for root physiology (Hirsch & Oldroyd, 2009). A nematode effector conserved in *Meloidogyne* spp. acts as a signalling molecule that specifically targets the plant SCL transcription regulators to induce root proliferation (Huang et al., 2006).Therefore, our results support the notion that the induction of SCL led to increased cell proliferation in the roots, which is essential for GCs induction and expansion. Further, we detected three nodulation-signaling pathway (NSP) genes 1 and 2 under positive regulation **(Fig 6D and S4 Table)**. In addition, we found that genes encoding nodulin-like proteins were either induced (7 genes) or repressed (5 genes) by nematode infection **(S5 Table)**. It has been reported previously that RKN can invoke similar host signals involved during the formation of nodules necessary for nitrogen fixation (Favery et al., 2002). In various phytopathosystems, nodulin-like genes are involved in solute supply during these interactions (Denancé et al., 2014). In the same way, nematode infection might have induced nodulin-like genes and their transcriptional regulators NSP1 and NSP2 in the GRAS family to aid in solute transportation in the GCs, a crucial process for a successful nematode infection. overall, our results show that potato susceptibility to RKN is controlled at the transcriptional level by a complex gene regulatory network.

### Nematode responsive phytohormones

Plant hormone signal transduction pathways are typically targeted by pathogens to either disrupt or avoid plant defense responses. Pathogen invasion results in changes in various plant hormone levels (Bari & Jones, 2009). In this context, our study shows that nematode infection influenced the expression of genes associated with the synthesis of JA, SA, ET, auxin, gibberellic acid (GA) and cytokinin (CK) signaling pathways **(Fig 7 and S5 Table).** Salicylic acid signaling pathway positively regulates immunity to biotrophic parasites while JA and ET hormones usually function synergistically to regulate defense against necrotrophic microbes and herbivorous insects (Bari & Jones, 2009). Differential expression of genes involved in the phenylpropanoid metabolic pathway was detected including three key enzymes, one gene encoding phenylalanine ammonia lyase (HAL), and one encoding trans-cinnamate (C4H, 3 genes) were repressed at 3 and 7dpi. Further, two genes encoding 4-coumarate-CoA ligase (4CL1) were repressed at 7 dpi and one gene slightly activated at 3dpi during RKN disease development **(S3 Table)**, which can affect other downstream activities such as SA, lignin and flavonoids biosynthesis (Vogt, 2010). Here, we detected regulation of genes in the flavonoid biosynthetic pathway, including chalcone synthase (induced, one gene), chalcone-flavone isomerase and flavanol sulfotransferase-like (repressed, one gene) at 3 and 7dpi **(S5 Table)**. Moreover, enzymes that participate in lignin biosynthesis were detected including cinnamyl alcohols dehydrogenase (CAD, one gene) and lignin-forming anionic peroxidase (five genes) were repressed in response to RKN. O-methyltransferase encoding genes in the lignin pathway were either slightly up-regulated or down-regulated at 3 and/or 7dpi **(S5 Table).** This demonstrates the involvement of phenylpropanoid metabolic pathway in RKN disease development.

**Figure 7:**
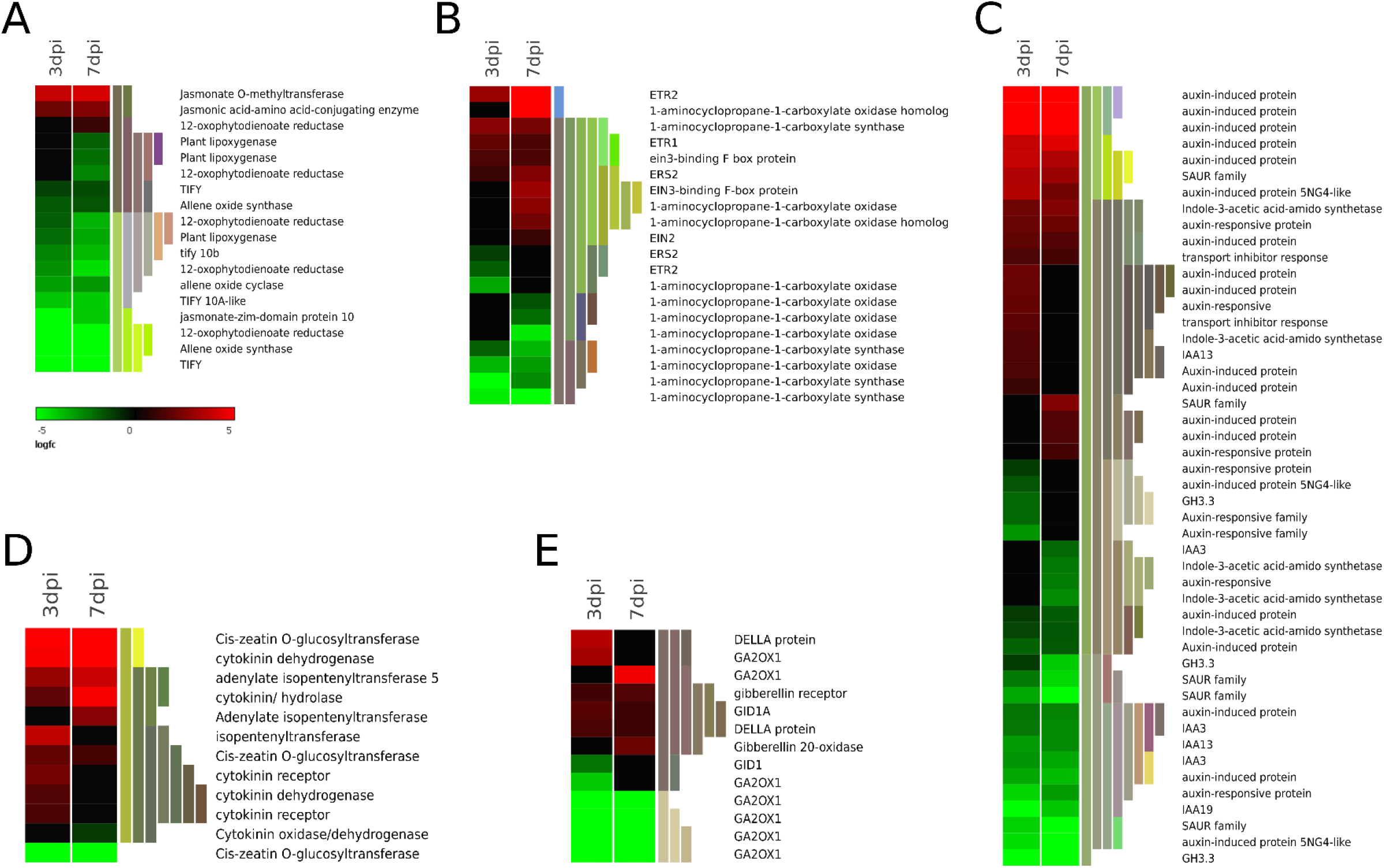
Heat map illustration of DEGs involved in hormone signal transduction. **(A)** JA signaling pathway. **(B)** ET signaling pathway. **(C)** Auxin signaling pathway. **(D)** Cytokinin signaling pathway. **(E)** GA signaling pathway. (The heat map illustrates a subset of genes from each group. Refer supporting information for all DEGs in each group).

For SA signalling, chorismate mutase (one gene) and SA-carboxyl methyltransferase (one gene) encoding genes implicated in SA synthesis (D’Maris Amick Dempsey et al., 2011) were down-regulated following nematode infection in addition to differential regulation of a gene encoding for key enzymes of PAL pathway **(S5 Table)**. Further, a subset of the WRKY family specifically involved in SA signalling pathway (i.e. WRKY70, WRKY40, WRKY17, and WRKY8) was suppressed according to our data set **(Fig 6B and S4 Table)**. These genes are important regulators of SA-dependent responses and have been implicated in the antagonistic crosstalk between SA-JA pathways (Pieterse et al., 2012) indicating repression of SA pathway by RKN infection.

Genes encoding for enzymes involved in the JA-signaling pathway were largely down-regulated. These include genes encoding allene oxide synthase (AOS 2 genes), allene oxide cyclase (AOC, one gene), lipoxygenase (LOX, 3 genes) and 12-oxophytodienoate (12-OPR, 5 genes) **(Fig 7A and S5 Table)**. The LOX pathway mediates resistance against pathogens, insects, and nematodes (Gao et al., 2008). Gleason et al. (2016) demonstrated that the 12-OPR enzyme, a JA-precursor, is a vital defense-signaling molecule that mediates plant immunity against nematodes. Moreover, plants incapable of producing JA or 12-oxo-phytodieonoic acid (OPDA) are more susceptible to phytonematodes (Gleason et al., 2016). In this perspective, the down-regulation of LOX and 12-OPR enzymes in the current study might have played a role in initiating a susceptible interaction through interfering with JA-mediated defense pathway. In addition, jasmonate O-methyltransferase, an additional regulatory point for the accumulation of jasmonate derivatives in the cytoplasm and production of signal transmitters other than JA (Seo et al., 2001) was up-regulated according to our dataset **(Fig 7A and S5Table)**. Interestingly, 5 genes of the TIFY protein family, which includes jasmonate-Zim-domain protein 10 (JAZ10), that represses JA signaling pathway, were down-regulated in this study **(Fig 7A and S5 Table)**. This indicates the activation of this pathway although not sufficient to mount the defense against RKN. Apart from pathogen and herbivory attack, the JA pathway can be activated as result of wounding. It also enhances accumulation of protease inhibitors which hinder exogenous proteases from insects to halt their development and reproduction (Koo & Howe, 2009). It is likely that JA-mediated defenses are effective against phytonematodes. However, the strong induction of several classes of protease inhibitors in our study **(S7 Table)** did not correlate with RKN resistance. Thus, the activation of the JA signaling pathway in this study might be due to wounding caused by nematode migration and feeding rather than by defense response.

Activation of the ET pathway upon pathogen attack leads to accumulation of defense-related through a cascade of events leading to activation of ERF TFs (van Loon et al., 2006). In the current data set, genes encoding for key enzymes involved in ethylene biosynthesis including 1-aminocyclopropane-1-carboxylate (ACC, 4 genes) synthase and 1-aminocyclopropane-1-carboxylate (ACO, 5 genes out of 8 genes) were down-regulated. We also detected three up-regulated membrane receptors (which perceive ET) at 3 and/or 7dpi **(Fig 7B and S5 Table)**. This includes ETHYLENE RESPONSE1 and 2 (ETR1 and ETR2) and ETHYLENE RESPONSE SENSOR2 (ERS2) negative regulators of ET responses (Ju & Chang, 2015) suggesting ET suppression. Further, ETHYLENE INSENSITIVE3 BINDING F-BOX (EBF) proteins (2 genes) that degrade EIN3/ETHYLENE INSENSITIVE3-LIKE1 (EIL) key positive regulators of ET responses (Ju & Chang, 2015) were activated in our data set **(Fig 7B and S5 Table)**. Our results show that apart from suppressing ET synthesis genes, RKN induced both negative regulators of ET responses and EBF responsible for proteasomal degradation of EIN3/EIL TFs that positively regulate ET responsive genes. This hindered the activation of defense-related genes associated with ERF branch of TFs.

Auxin stimulates several changes such as cell wall ingrowths, cell cycle activation and cell expansion occurring in nematode feeding sites (Gheysen & Mitchum, 2018). Here, we found 51 auxin signaling genes differentially expressed, including tryptophan aminotransferase-related protein 4 (auxin biosynthesis), GH3.3 and SAUR family (auxin-responsive genes), auxin repressors (e.g. IAA13, IAA19) and auxin transporters (e.g. TIR, LAX1) **(Fig 7C and S5 Table)**. It has been recently reported that nematode invasion results in induced auxin biosynthesis and responsive genes while genes encoding for repressors are switched off (Gheysen & Mitchum, 2018). Similar to this scenario, in our dataset, auxin repressors were repressed further highlighting the importance of auxin manipulation in nematode parasitism. Overall, *M. javanica* modulates auxin signaling pathway to facilitate successful formation of GCs.

Cytokinin and auxin hormones have been implicated in the induction and development of NFS (Gheysen & Mitchum, 2018). In the present study, genes involved in CK signaling pathway were differentially expressed. These include cytokinin dehydrogenase and cis-zeatin O-glucosyltransferase involved in CK homeostasis, of which 5 genes out of 7 were induced at 3 and/or 7 dpi **(Fig 7D and S5 Table).** Cytokinin dehydrogenase is involved in the degradation of CK. Transgenic plants overexpressing this enzyme had decreased gall formation and consequently reduced susceptibility to nematode infection (Lohar et al., 2004, Siddique et al., 2015). Due to their involvement in nutrient mobilization and cell division, CKs are believed to play a role in formation and maintenance of NFS infection (Lohar et al., 2004, Siddique et al., 2015). Our study shows that RKN regulates CK levels by regulating genes associated with the homeostasis of CK further underlining the significance of CK in GCs formation.

Next, our RNA-seq data revealed differential expression of the genes encoding for enzymes involved in GA biosynthesis including 3 up-regulated GA2OX1 encoding genes, and 5 down-regulated at both infection stages. Two genes encoding for DELLA proteins, which are negative regulators of GA response, were up-regulated while GA receptors were either up (2 genes) or down-regulated (one gene) by nematode infection at 3 and 7dpi **(Fig 7E, S5, and S4 Table)**. GA2OX1 enzymes reduce endogenous GA content in *Arabidopsis* plants that stimulates plant elongation process (Lee et al., 2014, Hu et al., 2017). A similar observation was made on tomato and rice plants, where GA2OX and GA receptors were strongly activated following RKN attack (Bar-Or et al., 2005, Kyndt et al., 2012). Further supporting this notion, GA foliar application on tomato plants enhanced resistance to *M. javanica* (Moosavi, 2017). Collectively, these results show that *M. javanica* modulates GA signaling process by activating GA20X1 enzymes and GA repressors, which reduce the active GA and stimulate root elongation that might be essential during GCs induction (Fuchs et al., 2013).

Several components of ABA stress-responsive hormone signaling, including ABA receptors, protein phosphatase 2C (PP2C) and SNF1-related protein kinases (SNRK) were differentially regulated in the present data set **(S5 Table)**. Genes encoding for PP2Cs were repressed (26 out of 38) by nematode infection according to our data set **(S3 Table)**. PP2Cs encoding genes are major players in stress signalling (Fuchs et al., 2013) that transmit ABA signaling directly from receptors to their downstream regulators. The SNRK regulators then activate an ABF/AREB/AB15 clade of bZIP-domain TFs through protein phosphorylation process finally to induce physiological ABA response(Sun et al., 2011). Apart from regulating stress responses and plant development, members of bZIP TF family are also implicated in plant defense response (Singh et al., 2002). In this study 23 bZIP genes were differentially expressed with 14 genes activated and 9 genes repressed **(Table S4)**. Therefore, we can hypothesize that nematode infection modulates the main stress-signaling pathway through repression of ABA receptors, which blocks the expression of some of the bZIP TFs responsible for defense response initiation.

### Genes associated with metabolic activities and transport activity are regulated by *M. javanica* infection

As obligate biotrophs, RKN fully depends on host-derived nutrients and solute transport to establish feeding sites. The differentiation of giant cells is coupled with massive changes in structure and metabolism of the host cells (Siddique & Grundler, 2015). GO enrichment analyses showed that genes involved in primary metabolism and cellular metabolism were overrepresented among the down-regulated genes **(S2 Fig and S1 Table).** Repression of these genes might be a strategy adopted by the host to save energy which is diverted for defense responses (Rojas et al., 2014). For instance, in this study, genes associated with lipid metabolism such as GDSL esterases/ lipases were repressed (14 genes) by nematode infection at both infection stages **(Fig 8A and S6 Table)**. Lipid and their metabolites have a role in mediating plant resistance (Gao et al., 2017). This indicates that *M. javanica* interfered with the host lipid-based defenses when initiating a compatible interaction. Nematode infection is associated with drastic reorganization of infected plant cells as well reprogramming of plant primary metabolism (Hofmann et al., 2010). It is also believed that nematodes may trigger biosynthesis of essential nutrients for their development, hence new metabolic pathways maybe induced in the host plants (Hofmann et al., 2010). Other genes in plant primary metabolism category under differential regulation of RKN parasitism include glycolytic process, trehalose metabolism, fatty acid biosynthetic process, sucrose and protein metabolism **(S6 Table)**.

**Figure 8:**
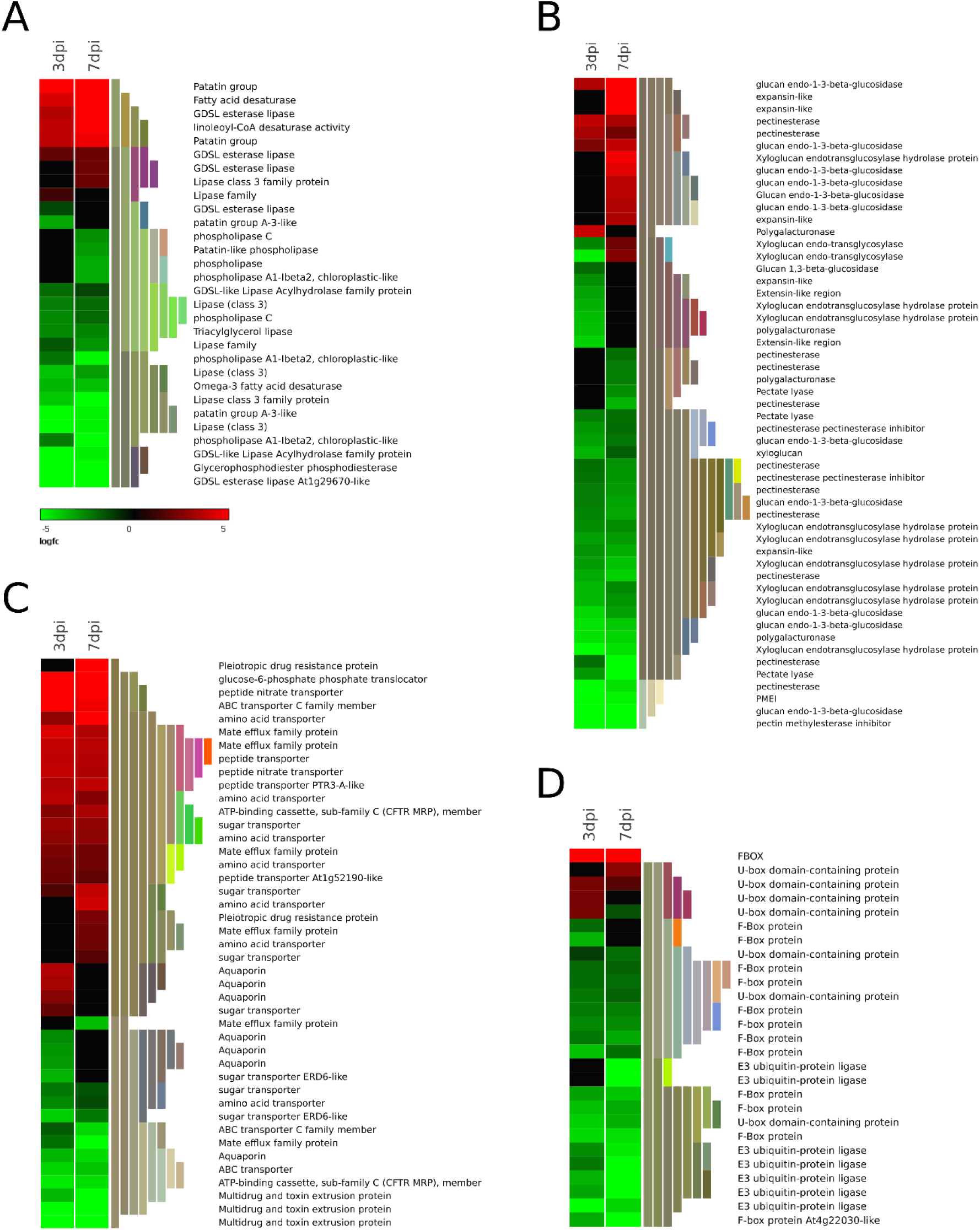
Heat map representation of gene expression patterns of genes associated with metabolism and transport activity **(A)** Lipid metabolism. **(B)** Cell wall architecture. **(C)** Transporters **(D)** Proteolysis and ubiquitination processes. (The heat map illustrates a subset of genes from each group. Refer supporting information for all DEGs in each group).

Similar to other studies, our transcriptomic data revealed that 165 genes encoding for cell wall modifying/degrading enzymes (CWM/DEs) annotated under the carbohydrate metabolic processes were differentially expressed. This includes genes encoding for glucan endo-1-3 beta-glucosidase, xyloglucan endo-transglycosylase, expansins, and extensins which were differentially expressed at 3 and/or 7dpi by RKN infection **(Fig 8B and S6 Table)** This shows that the regulation of these CWD/MEs is important during cell wall modification in the NFS. Glucan endo-1-3beta-glucosidase (members of PR-2 protein family) were differentially expressed with 14 genes down-regulated and 6 genes up-regulated at 3 and/7 dpi in this study **(Fig 8B and S6 Table)**. These cell wall modifying enzymes and also linked to plant defense against pathogens (van Loon et al., 2006). Callose deposition of a 1,3-ß glucan polymer is one of the first events occurring during pathogen invasion to slow down its progression (Voigt, 2014). Therefore, suppression of Glucan endo-1-3beta-glucosidase genes by nematode infection interferes with callose formation as well as plant defense. Among the 14 repressed genes encoding for xyloglucan endo-transglycosylase, two genes (PGSC0003DMG400004670 and PGSC0003DMG400021877) were induced by nematode infection at 7dpi (**Fig 8B and S6 Table**) suggesting a tight regulation of CWMEs during GCs formation. Similarly, 3 expansin encoding genes were specifically up-regulated at 7 dpi. **(Fig 8B and S6 Table)**. Shukla et al. (2018) also demonstrated the up-regulation of genes encoding for expansins between 5 and 7 dpi in a susceptible tomato response to RKN. Expansins are cell wall loosening proteins that might play key role during the expansion of GCs (Gheysen & Mitchum, 2008). In addition, genes encoding for hydrolytic enzymes involved in pectin degradation such as polygalacturonase (PG), pectate lyases (PL) and pectin esterase (PE) were differentially expressed at 3 and 7 dpi by nematode infection **(Fig 8B and S6 Table).** Pectin degradation leads to plant tissue maceration that is essential in disease development (Lionetti et al., 2012).Therefore, the reduction of pectin content may increase nutrient accessibility to nematodes (Jammes et al., 2005). Furthermore, four genes encoding pectin methyl esterase inhibitor (PMEI or PEI) were down-regulated upon nematode challenge at 3 and 7 dpi **(Fig 8B and S6 Table)**. Plants produce PMEI in an effort to counteract the increasing pectin methyl esterase (PME) upon pathogen attack (Lionetti et al., 2012). Repression of PMEI by nematode attack shows that activity of PME was activated leading to the breakdown of pectin bonds, which increases the vulnerability of the cell wall to microbial pectic enzymes and other degrading enzymes and culminates in a susceptible response. Generally, the differential regulation of genes associated with cell wall architecture suggests that *M. javanica* was able to break down plant cell wall to facilitate migration and formation of GCs. Furthermore, RKN interferes with defense role associated with the plant cell wall architecture leading to compatible interaction.

With the increased demand for nutrients in nematode feeding cells, nematodes deploy specialized membrane transporters to control the flow of nutrients in and out of the NFS (Rodiuc et al., 2014, Siddique & Grundler, 2015). In agreement with previous studies (Hammes et al., 2005, Shukla et al., 2018), we found that several families of transporter genes were differentially regulated upon RKN infestation in our analyses. These include peptide transporters, aquaporins, amino acid transporters, ion transporters, sugar transporters and glutathione S transferase **(Fig 8C and S6 Table)**. Overall, we found that 54.8% of transporter encoding genes in the DEGs were up-regulated following nematode infection. The activation of genes encoding amino acid transporters (8 genes) and sugar transporters (9 genes) indicates activation of amino acid and carbohydrate metabolism, respectively. For instance, according to Zhao et al. (2018), the induction of sugar transporters increases soluble sugar contents in RKN infected tomato plants, which is crucial for nematode development. Furthermore, multidrug transporter-encoding genes were differentially expressed in our samples as well **(Fig 8C and S6 Table),** which encompasses ATP-binding cassette (ABC, 16 genes) and multidrug and toxin extrusion proteins (MATE, 17 genes). These are secondary active transporters involved in plant immunity and transporting and trafficking of xenobiotic, small organic molecules, and secondary metabolites (Peng et al., 2011). In a similar study involving tomato and *M. incognita,* 15 MATE efflux proteins exhibited differential expression (Shukla et al., 2018), nevertheless, their role in plant-nematode interaction is yet to be defined. Hence, it is likely that nematodes recruit some of these transporters to flush out toxic secondary metabolites or to disperse nematode effectors produced following nematode invasion.

Nematode effectors that target plant ubiquitin-proteasome system (UPS) have been reported previously (Rehman et al., 2009, Chronis et al., 2013). In the present study, we found that several genes involved in protein ubiquitination and proteolysis including the U-box domain (27 genes) RING finger protein (3 genes) zinc finger domain (8) and the F-box (30 genes) were down-regulated **(Fig 8D and S7 Table)**. This could indicate an immense turnover of proteins due to constant nematode feeding leading to a compatible interaction. Wang et al. (2015) reported an enzyme E3 ligase, U-box/ARM repeat protein (OsPUB15) which interacts with a receptor-like kinase to regulate programmed cell death as well as disease resistance. Similarly, E3 ubiquitin ligase protein is known to control plant immunity to a broad range of microbes in rice through orchestrating plant immunity homeostasis and coordinating the trade-off between defense and growth in plants (You et al., 2016). Indeed, the findings further support that the UPS system might be a primary target to *M. javanica* effectors, which modulate the various facets of plant defense linked with the UPS system.

Collectively, this study uncovers the molecular networks regulated during compatible interaction between potato and RKN. This provides further insights on plant-nematode interactions and will enhance further studies in this area including development of target-specific control strategies against *Meloidogyne* species.

## Materials and methods

### Plant material and RKN inoculations

Certified seed (tubers) of seven potato cultivars were grown under greenhouse conditions to evaluate resistance to *M. javanica* under greenhouse conditions. The seed tubers were pre-germinated in the dark 20 ± 3°C for seven days to allow sprouting. Stocks of *M. javanica* were originally obtained from Dr. Pofu (ARC Roodeplaat, Pretoria, South Africa) and maintained on susceptible tomato cultivar, *S. lycorpersicum* Cv Floradade in glasshouse environment with a temperature of 24 °C-30 °C for eight weeks. *Meloidogyne javanica* eggs were extracted from infected roots as described (Hussey, 1973). Egg suspension was poured onto an extraction tray for collection of second juveniles’ (J2s) nematodes. Five-week old potato seedlings were inoculated with 1000 freshly hatched J2s per plant and control plant mock-inoculated with water. The number of galls using Taylor and Sasser (1978) ranking scale to determine susceptibility and reproduction factor (RF) using Sasser et al. (1984) RF formula was used to assess the host status of potato cultivars to RKN infection 8 weeks after infection For RNA experiment, whole root tissues of a compatible potato cultivar were collected at 0, 3 and 7 days post-inoculation (dpi) with two biological replicates per time point. Samples were washed and immediately frozen in liquid nitrogen to prevent RNA degradation and later stored at −80 °C until RNA extraction.

### RNA extraction, library preparation, and sequencing

RNA extraction, library preparation, and sequencing were carried out at Novogene (HK) Company Limited. Total RNA for individual time course and replicates was extracted using TiaGen extraction kit (Biotech Beijing Co., Ltd) and treated with sigma DNase1 (D5025). RNA degradation and contamination was measured on 1% agarose gel while RNA purity was assessed using the NanoPhotometer^®^ spectrophotometer (IMPLEN, CA, USA). RNA concentration and integrity were assessed using Qubit ^®^ RNA Assay kit in Qubit ®2.0 Fluorometer (Life Technologies, CA, USA) and RNA Nano 6000 Assay Kit of the Bioanalyzer 2100 system (Agilent Technologies, CA, USA), respectively. Three micrograms of RNA samples were used as input for library construction. Libraries were constructed using NEBNext® Ultra™ RNA Library Prep Kit for Illumina^®^ (NEB, USA) according to the manufacturer’s instructions and index codes were added to attribute sequences to each sample. Finally, PCR products were purified using AMPure XP system and quality of the library assessed using the Agilent Bioanalyzer 2100 system. A cBot Cluster Generation System using HiSeq PE Cluster Kit cBot-HS (Illumina) was used to cluster the index-coded samples. After cluster generation, the library preparations were sequenced on an Illumina Hiseq platform 2500 generating 150 bp paired-end reads.

### Transcriptomic data analysis

Quality analysis of sequenced reads were initially analyzed using FASTQC package (https://www.bioinformatics.babraham.ac.uk/projects/fastqc). Clean reads were obtained by removing reads containing adapter reads with poly-N and low-quality reads from raw data. Trimming of low-quality regions was performed using Trimmomatic v 0.36 (Bolger et al., 2014). All the subsequent downstream analyses were based on high-quality data. *Solanum tuberosum* genome v4.03 (Consortium, 2011) was used for reference-guided mapping of RNA-seq reads. Paired-end clean reads were aligned to the potato genome using hisat2 v 2.1.0 software (Kim et al., 2015). Unmapped reads were progressively trimmed at the 3’end and re-mapped to the genome. Next, featureCounts package (Liao et al., 2014) was used to perform raw-reads counts in R environment (https://www.r-project.org/). The read counts were then used for differential expression analysis using edgeR package (Robinson et al., 2010). Further, to investigate the responses at different time points (3 dpi and 7 dpi), the expression profiles were compared to mock-inoculated (0dpi) data sets. The transcripts were then classified as differentially expressed genes (DEGs) based on both (a) false discovery rate (FDR) (Benjamini & Yekutieli, 2005) cut-off of 0.05 and (b) log2 fold change ≥ 1 or ≤ −1 for induced and repressed genes, respectively.

### Gene ontology (GO) and enrichment analysis

The GO and enrichment analysis were performed using agriGO v.2.0 (Tian et al., 2017) and categorized by WEGO v 2.0 tool (Ye et al., 2018). Parametric gene set enrichment analysis based on differential expression levels (log2 fold change) was performed and FDR correction was performed using the default parameters to adjust the *p*-value. Functional annotations and pathway analyses were obtained through sequence search performed on eggNOG database utilizing *eggmapper* (Huerta-Cepas et al., 2017). Annotations from eggNOG were then integrated with Kyoto Encyclopedia of Genes and Genomes (KEGG) database in order to reach pathway annotation level.

### Validation for DEGs by qRT-PCR

For qRT-PCR, first-strand cDNA was done from total RNA using Superscript IV First-Strand cDNA Synthesis SuperMix Kit (Invitrogen, USA) following manufacturer’s protocol. Quantitative real-time PCR was performed using SYBR Green Master Mix in the QuantStudio 12k Flex Real-Time PCR system (Life Technologies, Carlsbad, CA, USA) to validate DEGs. Two micrograms of the sample was added to 5 μl of Applied Biosystems SYBR Green Master Mix and primers at a concentration of 0.4 μM. The implication cycle consisted of following: initial denaturation at 50 °C for 5 min and 95 °C for 2 min followed by 45 cycles of 95 °C for 15 s and 60 °C for 1 min. Each sample was run in triplicates. Specific qRT-PCR primers for six target genes were designed using an online tool Prime-Blast (http://www.ncbi.nlm.nih.gov/tools/primer-blast) **(S8 Table).** Each sample was run in triplicates. The 18S rRNA and elongation factor 1-α (PGSC0003DMG400020772,ef1α), (Nicot et al., 2005) were used as the reference genes for normalization and the mock-treated samples used as calibrators. The comparative 2^-ΔΔ*Ct*^ method was used to determine the relative fold change according to Schmittgen and Livak (2008). Despite, the two techniques (RNA-seq and qRT-PCR) being different, the expression patterns of selected genes upon nematode infection was consistent between the two procedures **(Fig S9)**.

### Data access

Both raw and processed sequencing data have been deposited to the Gene Expression Omnibus (GEO) repository at the National Center for Biotechnology Information (NCBI) with accession no. GSE134790.

## Acknowledgments

This research study was funded by the National Research Foundation (NRF), South Africa through Competitive Funding for Rated Researchers (CFRR) 98993, Bioinformatics and Functional Genomics (BFG 93685) and Potatoes South Africa (PSA). DB-R was supported by University of Pretoria Post-Doctoral Fellowship. TM was funded by Potato South Africa and the NRF Scarce Skills/Innovation Scholarships.

## Author’s Contributions

L.N.M conceived, designed this study and funding acquisition. T.N.M set-up the experiment for nematode inoculations, analyzed and visualized data and wrote the original draft. D.R.B carried out the bioinformatics work. L.N.M and D.R.B revised the manuscript. All authors reviewed and made changes to the initial draft and approved the final version.

## Abbreviations

CK: Cytokinin
DEGs: Differentially expressed genes
ET: Ethylene
GO: Gene ontology
GCs: Giant cells
GA: Gibberellic acid
JA: Jasmonic acid
NFSs: Nematode feeding sites
PR: Pathogenesis-related protein
PTI: Pattern triggered immunity
ROS: Reactive oxygen species
RKN: Root-knot nematode
SA: Salicylic acid
TFs: Transcription Factors

