## Supplementary Figure 1 for "Transcriptional profiling of potato (*Solanum tuberosum* L.) during a compatible interaction with the root-knot nematode, *Meloidogyne javanica*"

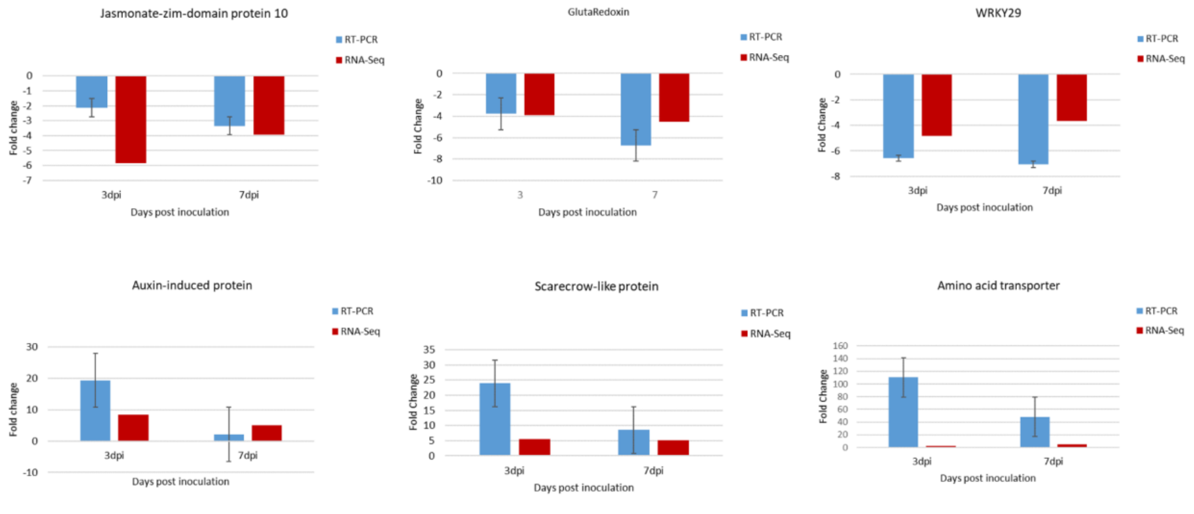


**Figure S1.** The expression profiles of eight genes experimentally validated using real-time-qPCR in response to nematode invasion at 3 and 7dpi. The qPCR error bars represent the standard error from three biological replicates.
